# Structural Dissection of Vaccinia G9 Identifies Residues Essential for Membrane Fusion and Complex Assembly

**DOI:** 10.1101/2025.04.22.650039

**Authors:** Hsiao-Jung Chiu, Hao-Ching Wang, Wen Chang

## Abstract

Vaccinia virus, a prototypical poxvirus, utilizes a unique multi-protein Entry Fusion Complex (EFC), comprising 11 components, to mediate membrane fusion during host cell entry. Although the crystal structure of a truncated form of the G9 protein has been determined, the functional relevance of its structural features remains poorly understood. In this study, we systematically analyzed 47 G9 mutants to identify critical functional residues. Using trans-complementation assays, co-immunoprecipitation, membrane fusion assays, and structural analysis, we identified nine key mutants, which were categorized into three functional groups. Group 1 mutants failed to interact with A16 and other EFC components, highlighting their essential roles in G9-A16 subcomplex formation. Group 2 and Group 3 mutants retained A16 binding but disrupted interactions with other EFC proteins, suggesting their roles in broader complex assembly. Notably, Group 3 mutants targeted a conserved P(R/Y)XCW motif and a loop structure shared among vaccinia G9, A16, and J5 proteins. A similar motif was also identified in G9 homologs from *Nucleocytoviricota*, suggesting an evolutionarily conserved fusion mechanism. Collectively, our findings demonstrated that G9 function requires multiple domains, including A16-binding interfaces and conserved motifs not resolved in previous protein structure. These results establish G9 as a central EFC component and underscore its potential as a target for antiviral development.

**Importance:** Understanding how viruses enter host cells is critical for developing antiviral strategies. Vaccinia virus, a model poxvirus, uses a unique 11-protein entry fusion complex (EFC) to mediate membrane fusion—unlike other viruses that rely on a single fusion protein. In this study we identified specific residues in the G9 protein that are critical for maintaining EFC function. Notably, we discovered a conserved P(R/Y)XCW motif within G9 that is also present in orthologs from both poxviruses and members of the *Nucleocytoviricota* phylum, suggesting an evolutionarily conserved mechanism of membrane fusion. These conserved structural elements can serve as potential targets for antiviral intervention against pathogenic poxvirus infections in humans.

## Introduction

Vaccinia virus is a large double-stranded DNA virus belonging to the genus *Orthopoxvirus* in the *Poxviridae* family (1). Its genome is approximately 190 kbp in length and encodes over 200 proteins (2). Vaccinia virus contains two infectious forms of virus particles produced from the infected cells, the mature virus (MV) and the enveloped virus (EV). The majority of the virus particles (>95%) accumulated intracellular as MV, which are brick-shaped with membranes derived from the endoplasmic reticulum membranes (3, 4). A small fraction of MV (∼5%) are transported to the Golgi apparatus, where they acquire two additional membrane layers derived from the Golgi cisternae, forming wrapped virions (WVs) (5, 6) These WVs are transported along microtubules to the cell periphery (7, 8), where they fuse in an inside-out manner with the plasma membrane, releasing extracellular EV (9, 10).

Vaccinia MV contains four attachment proteins- H3 (11), D8 (12), A27 (13), and A26 (14). Membrane fusion is mediated by an 11-protein Entry Fusion Complex (EFC), composed of G9 (15), A16 (16), J5 (17), A21 (18), A28 (19), H2 (20), G3 (21), L5 (22), F9 (23), L1 (24), and O3 (25), all of which are highly conserved across the *Poxviridae* family (26). Deletion or inhibition of any individual EFC component resulted in defective membrane fusion, demonstrating that each is essential for EFC function (26). Although the EFC becomes destabilized when specific components are lost, subunit interactions such as G9-A16, H2-A28, and G3-L5 remain stable (27–29). Recent studies suggested that L1 and F9 interact weakly with the rest of the complex, indicating they may serve as the peripheral rather than core components (30). Notably, while individual EFC proteins contribute to distinct stages of membrane fusion process, G9, A16, A21, H2, G3, F9, and O3 appear to function in the initiation of hemifusion, whereas A28, L1, and L5 act in post-hemifusion steps (31). Although the ectodomain structures of nearly all EFC components, except J5 and O3, have been resolved (32–38), the molecular mechanism of EFC-mediated membrane fusion remains poorly understood.

To regulate membrane fusion, vaccinia virus encodes fusion suppressors that keep the activity of the Entry Fusion Complex (EFC) under control. One such suppressor, the A26 protein, directly binds to the G9-A16 subcomplex and inhibits premature fusion activation at neutral pH (39). Another suppressor complex, composed of A56 and K2, is expressed on the surface of infected cells and also interacts with the G9-A16 subcomplex to prevent superinfection (30). Interestingly, experimental evolutionary studies have shown that a single H44Y mutation in the G9 protein can overcome A56/K2-mediated inhibition, enabling membrane fusion to occur at pH 6 instead of pH 5 (30). Furthermore, phylogenetic analyses revealed that G9 and A16 homologues of the EFC are present in *Nucleocytoviricota*, implying that the vaccinia EFC evolved from a shared ancestral fusion machinery (40). In addition, phylogenetic analyses revealed that homologs of G9 and A16 are present in members of the phylum *Nucleocytoviricota*, suggesting that the vaccinia EFC may have evolved from an ancestral fusion apparatus shared among large DNA viruses. Given the essential role of G9 protein in EFC-mediated membrane fusion, we sought to dissect its structural and functional relationship. Using site-directed mutagenesis, we identified critical regions of G9 required for viral infectivity, subcomplex formation and assembly of an intact EFC. These findings provide structural insights into the contribution of G9 protein to membrane fusion in vaccinia virus.

## Material and methods

### Cell culture and virus

Human 293T and African green monkey kidney BSC40 cells were maintained in Dulbecco’s Modified Eagle Medium (DMEM) supplemented with 10% fetal bovine serum (FBS)(36). Recombinant vaccinia virus vG9Li-HA, which expresses the vaccinia G9 protein under induction with isopropyl-β-d-thiogalactopyranoside (IPTG), was kindly provided by B. Moss (15). The vG9Li-HA virus was propagated in BSC40 cells cultured in medium supplemented with 50 µM IPTG, as previously described (15).

### Construction of G9 expression plasmids by site-directed mutagenesis

The vaccinia *G9R* gene, tagged at the N-terminus with a c-myc epitope, was cloned into mammalian expression vectors pCAGEN or pcDNA3.1 to generate pCAGEN-myc-G9R or pcDNA3.1-myc-G9R. Mutant G9 constructs were generated using the QuikChange Lightning Site-Directed Mutagenesis Kit (Agilent) based on either of the two plasmids. All constructs were sequence-verified (Genomics Inc., Taiwan).

### Trans-complementation assays

Trans-complementation assays were performed as previously described (36). Briefly, 293T cells were seeded in 6-well plates and transfected with 1 µg of wild-type (WT) or mutant G9 plasmid using 10 µL Lipofectamine 2000 (Invitrogen). After 24 h, the cells were infected with vG9Li-HA at a multiplicity of infection (MOI) of 1 PFU/cell at 37°C for 1 h, followed by incubation in growth medium without IPTG. At 24 h post-infection (hpi), cells were harvested for immunoblot and plaque assays. Each mutant was tested in three independent experiments unless otherwise stated.

### Co-immunoprecipitation of G9 and EFC components

BSC40 cells in 100-mm dishes were infected with vG9Li-HA at an MOI of 5 PFU/cell for 1 h at 37°C, followed by transfection with 0.5 µg of WT or mutant G9 plasmid using 20 µL Lipofectamine 2000 in IPTG-free medium. At 24 hpi, cells were lysed on ice in 500 µL of lysis buffer (0.5% NP-40, 20 mM Tris-HCl, pH 8.0, 200 mM NaCl) supplemented with protease inhibitors (2 µg/mL aprotinin, 1 µg/mL leupeptin, 0.7 µg/mL pepstatin, 1 mM PMSF). Lysates were clarified by centrifugation at 16,000 × g for 15 min. Supernatants were incubated with c-myc-conjugated agarose beads (Sigma-Aldrich) at 4°C overnight and analyzed by immunoblotting with anti-EFC component antibodies (36). Each experiment was independently performed twice.

### MV-triggered cell-cell fusion at acidic pH

Cell-cell fusion assays were conducted as described previously (36). HeLa-RFP and HeLa-GFP cells were co-seeded in 96-well plates at a 1:1 ratio and treated with 40 µg/mL cordycepin for 1 h at 37°C before infection. WT or mutant G9 virus-containing lysates (from trans-complementation assays) were used to infect cells for 1 h at 37°C. Cells were then washed and exposed to pH 7.0 or pH 5.0 buffer at 37°C for 3 min, neutralized with growth medium, and incubated with cordycepin-containing medium for 3 h. Images were acquired from 30 min to 3 hpi using an ImageXpress Confocal HT.ai High Content Imaging System.

Fusion efficiency was quantified using Fiji software as: (% fusion) = (area of GFP⁺RFP⁺ cells / area of single-positive cells) × 100%, as previously described (35). WT G9-triggered fusion at low pH was set at 100%, and values for each mutant were normalized accordingly.

### Electron microscopy (EM)

BSC40 cells were infected with vG9Li-HA at an MOI of 5 PFU per cell and transfected with 1 µg of WT or mutant G9 plasmid using 10 µL Lipofectamine 2000 (Invitrogen). At 24 hpi, cells were fixed with glutaraldehyde and followed by secondary fixation with osmium tetroxide. The samples were stained with uranyl acetate, dehydrated, and embedded in Epon resin for sectioning as described (39). Sample sections were imaged using a Tecnai G2 Spirit TWIN transmission electron microscope.

### The multiple sequence alignments of G9 orthologues in *Poxviridae*

Orthologs of vaccinia G9 protein from various *Poxviridae* members (e.g., VACV, CMLV, CPXV, ECTV, MPXV, etc.) were retrieved from GenBank. Accession numbers and virus abbreviations are listed: vaccinia virus(YP_232969.1); CMLV: camelpox virus (NP_570475.1); CPXV: cowpox virus (NP_619884.1); ECTV: ectromelia virus (NP_671589.1); MPXV: monkeypox virus (NP_536506.1); SKPV: skunkpox virus (YP_009282780.1); TATV: taterapox virus (YP_717397.1); VARV: variola virus (NP_042116.1); VPXV: volepox virus (YP_009281834.1); EWPV: fowlpox virus (NP_039090.1); TKPV: turkeypox virus (YP_009177111.1); GTPV: goatpox virus (YP_001293250.1); YKPV: yokapox virus (YP_004821420.1); CRPV: crocodilepox virus (QGT49326.1); SQPV: squirrelpox virus (YP_008658478.1); MYXV: myxoma virus (NP_051768.1); MCPV: molluscum contagiosum virus (NP_044019.1); SEPV: sea otter poxvirus (YP_009480597.1); BPSV: bovine papular stomatitis virus (NP_957955.1); RDPV: red deer parapoxvirus (YP_009112785.1); PCPV: pseudocowpox virus (YP_003457351.1); ORFV: orf virus (QLI57553); PTPV: pteropox virus (YP_009268779.1); SGPV: salmon gill poxvirus (AKR04265.1); YMTV: yaba monkey tumor virus (NP_938314.1); ACEV: anomala cuprea entomopoxvirus (YP_009001529.1); EPTV: eptesipox virus (YP_009408014.1); and AMEV: amsacta moorei entomopoxvirus (NP_064817.1). Multiple sequence alignments were generated using MAFFT v7.453 with parameters --maxiterate 1000 --globalpair, and visualized with Jalview v1.8.3. Residue conservation was color-coded: yellow (identical), blue (>0.5 conservation), and green (>0.2 conservation).

### Protein Structure Prediction with AlphaFold2

Full-length G9 (YP_232969.1), A16 (YP_233018.1), and J5 (YP_232979.1) structures were predicted using AlphaFold2 (https://reurl.cc/96anZj). Sequences were queried using MMseqs2 (mmseqs2_uniref_env) in unpaired/paired mode. The predicted model with a pLDDT score >70 was selected for each protein. Final confidence scores for G9, A16, J5, and the G9-A16 subcomplex were 86.9, 81.2, 80.7, and 78.3, respectively.

### Residue Interaction Analysis

The G9-A16 subcomplex structure (PDB: 8GP6) (38) was analyzed using the Residue Interaction Network Generator (RING; https://ring.biocomputingup.it/) to identify intra- and inter-subunit residue contacts.

### Statistical analysis

Statistical analyses were performed using Student’s *t-test* in GraphPad Prism v.9.2. Data are presented as mean ± standard deviation (SD).

Comparisons with WT control were considered significant at: \**p* < 0.05, \*\**p*< 0.01, and \*\*\**p* < 0.001, and **** *p* < 0.0001.

## Results

### Identifying functional region of VACV G9 protein

To define the functional role of the vaccinia G9 protein during infection, we performed systematic mutagenesis of the G9 open reading frame (ORF), targeting two categories of residues: (i) surface-exposed charged residues (type I) and (ii) conserved residues among G9 orthologs in the *Poxviridae* (type II), as shown in Figure 1A. A total of 47 mutant constructs were generated and evaluated for their ability to support virus replication using a trans-complementation assay in 293T cells. Plasmids expressing either wild-type (WT) or mutant G9 proteins were transfected into 293T cells 24 hours prior to infection with the IPTG-inducible virus vG9Li-HA (15) at an MOI of 1 PFU/cell in the absence of IPTG. This setup ensures that G9 protein was only expressed from the transfected plasmid. At 24 hpi, cells were harvested for plaque assays and immunoblot analyses. All 47 mutant proteins were successfully expressed (Immunoblots in Figures 3A-C, 4A&B and 5E&F). As expected, WT G9 fully complemented the vG9Li-HA virus, yielding 100% infectivity (Figure 1B, black bar). In contrast, mutant constructs exhibited a wide range of phenotypes: eight mutants retained 50-75% infectivity (Figure 1B, yellow bars), three showed 25-50% infectivity (Figure 1B, orange bars), and nine had infectivity below 25% (Figure 1B, red bars). The last group of nine low-infectivity mutants were selected for further functional characterization.

**Figure 1.**
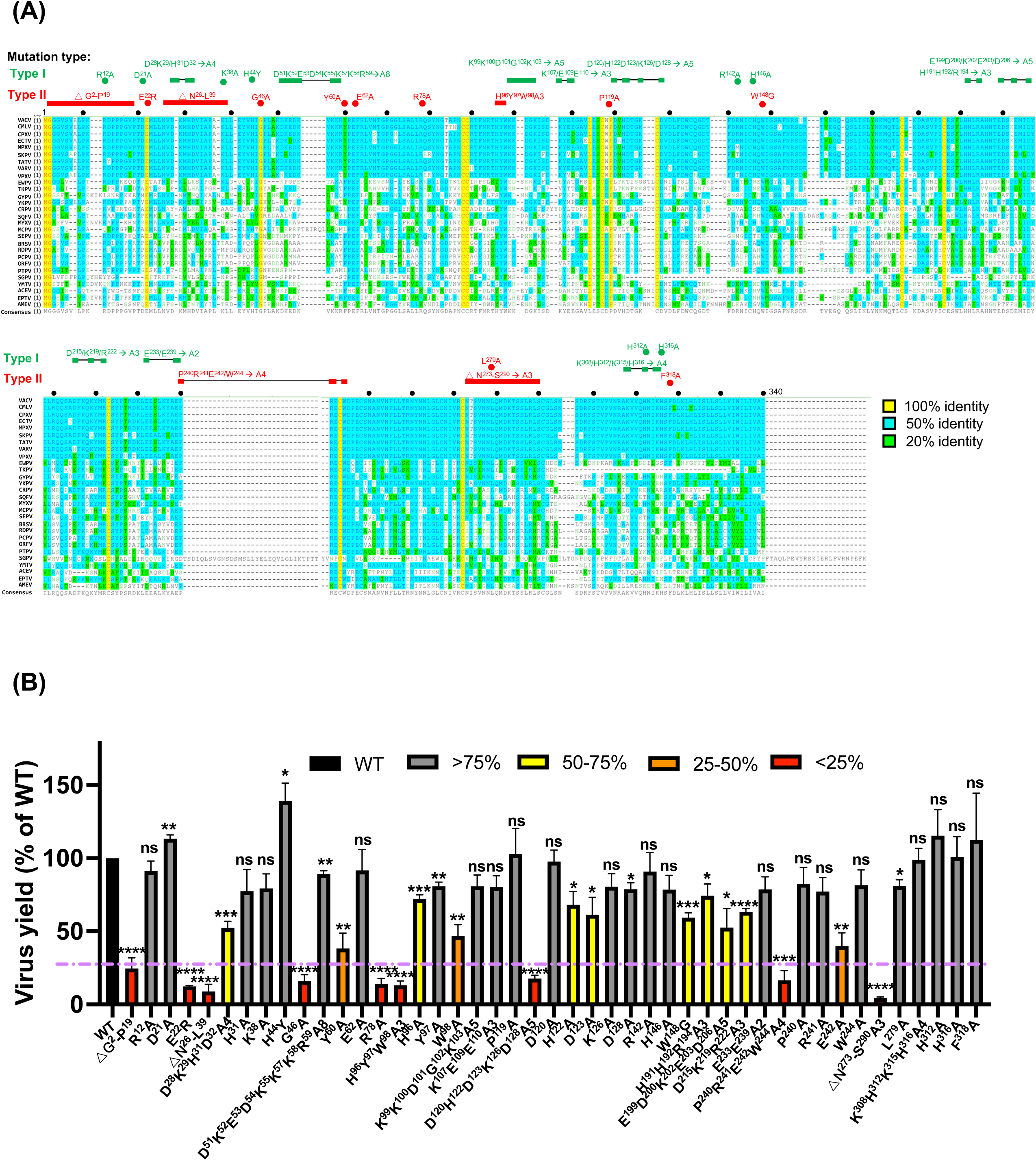
Identification of G9 Critical Regions important for vaccinia MV Infectivity. **(A)** Two types of G9 mutations were depicted on top of the Multiple sequence alignment of 28 G9 orthologues in *Poxviridae*: Mutations of the charged surface residues (type I) and conserved residues (type II) are shown in green and red, respectively. Colored dots and bridged-boxes represent mutation sites. Black dots mark the distance of 10 a.a. in the MSA. **(B)** Virus yields of G9 mutants from Trans-complementation assays. 293T cells were transfected with wild-type or mutant G9 plasmid and infected with vG9Li-HA and cultured for 24h without IPTG and harvested at 24h p.i. for plaque assays. The percentage of virus yield in each G9 mutant was normalized with that of WT G9. The purple dashed line indicates the 25% infectivity threshold. Student’s *t*-test: \**P* < 0.05, \*\**P* < 0.01, \*\*\**P* < 0.001, \*\*\*\**P* < 0.0001.

### G9 mutants with low MV infectivity are defective in membrane fusion

Previous studies using fluorescence dequenching analyses showed that G9-deficient vaccinia MV particles failed to initiate hemifusion with host membranes (31). To explore whether the critical residues we identified above are also required for fusion activity, we performed MV-triggered cell-cell fusion assays using virus-containing lysates from the trans-complementation experiments as previously described (35, 36, 39). It is known that repression of vaccinia EFC gene expression did not impair virion morphogenesis and MV particles of normal morphology were produced from the infected cells ((36) and references therein). Consistently, our G9 mutants did not impair MV morphogenesis (Fig. S1), validating the use of these inf/tf lysates for MV-triggered cell fusion assays. HeLa cells expressing GFP or RFP were mixed at a 1:1 ratio and infected with WT or mutant G9 virus lysates, treated with neutral pH 7 or acidic pH 5 buffer, and MV-triggered cell-cell fusion was monitored. As expected, no cell-cell fusion was triggered by WT nor mutant G9 viruses at neutral pH (Figure 2A, top panel). As expected, WT G9 virus triggered robust cell-cell fusion under acidic condition while none of the nine G9 mutants triggered detectable fusion (Figure 2A, bottom panel). Quantification of MV-triggered cell fusion at the low pH (Figure 2B) revealed that all nine G9 mutants lacked membrane fusion activity, indicating these residues are essential for EFC-mediated membrane fusion.

**Figure 2.**
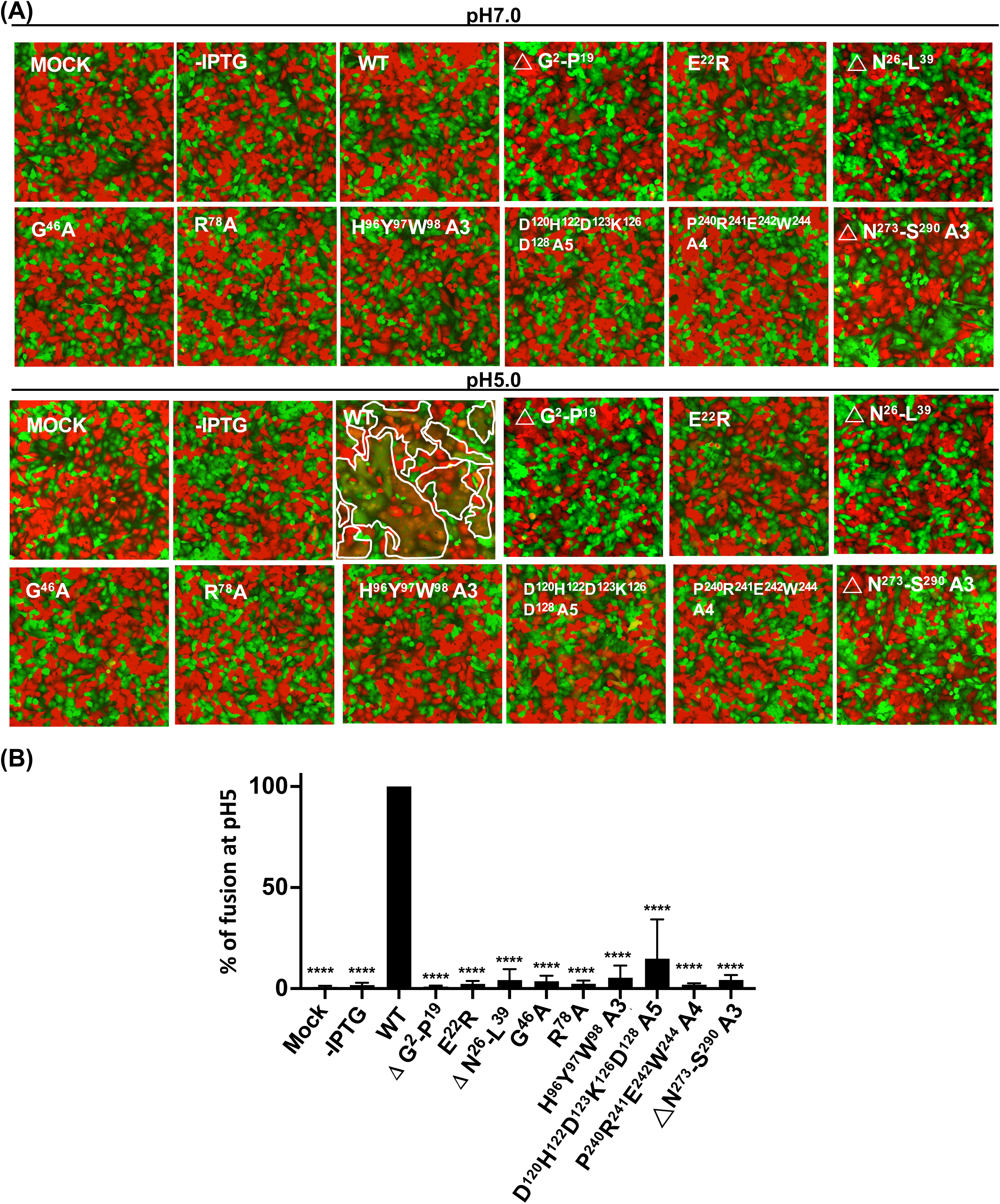
Cell-cell fusion assays of G9 mutants revealed impaired fusion at low pH. **(A)** HeLa cells expressing GFP or RFP were mixed at a ratio of 1:1 and treated with lysates containing WT or mutant G9 viruses in neutral pH 7 (top panel) or acidic pH 5 buffer (bottom panel). Fluorescent cell images were photographed at 3 hpi. The areas marked by white lines represented the fused cells. **(B)** Quantification of MV-induced cell-cell fusion at pH 7 and pH 5 from the images in (A). Photographs from three independent experiments were analyzed using Fiji software and the percentages of virus-triggered cell fusion at pH 5 were calculated as (the surface area of GFP^+^RFP^+^ double-fluorescent cells divided by the surface area of single-fluorescent cells) x100%. Error bars denote standard deviations from three independent experiments. Statistical comparisons of cell-cell fusion are between WT and each mutant at the low pH of 5. Student’s *t*-test: \**P* < 0.05, \*\**P* < 0.01, \*\*\**P* < 0.001, \*\*\*\**P* < 0.0001.

**Figure 3.**
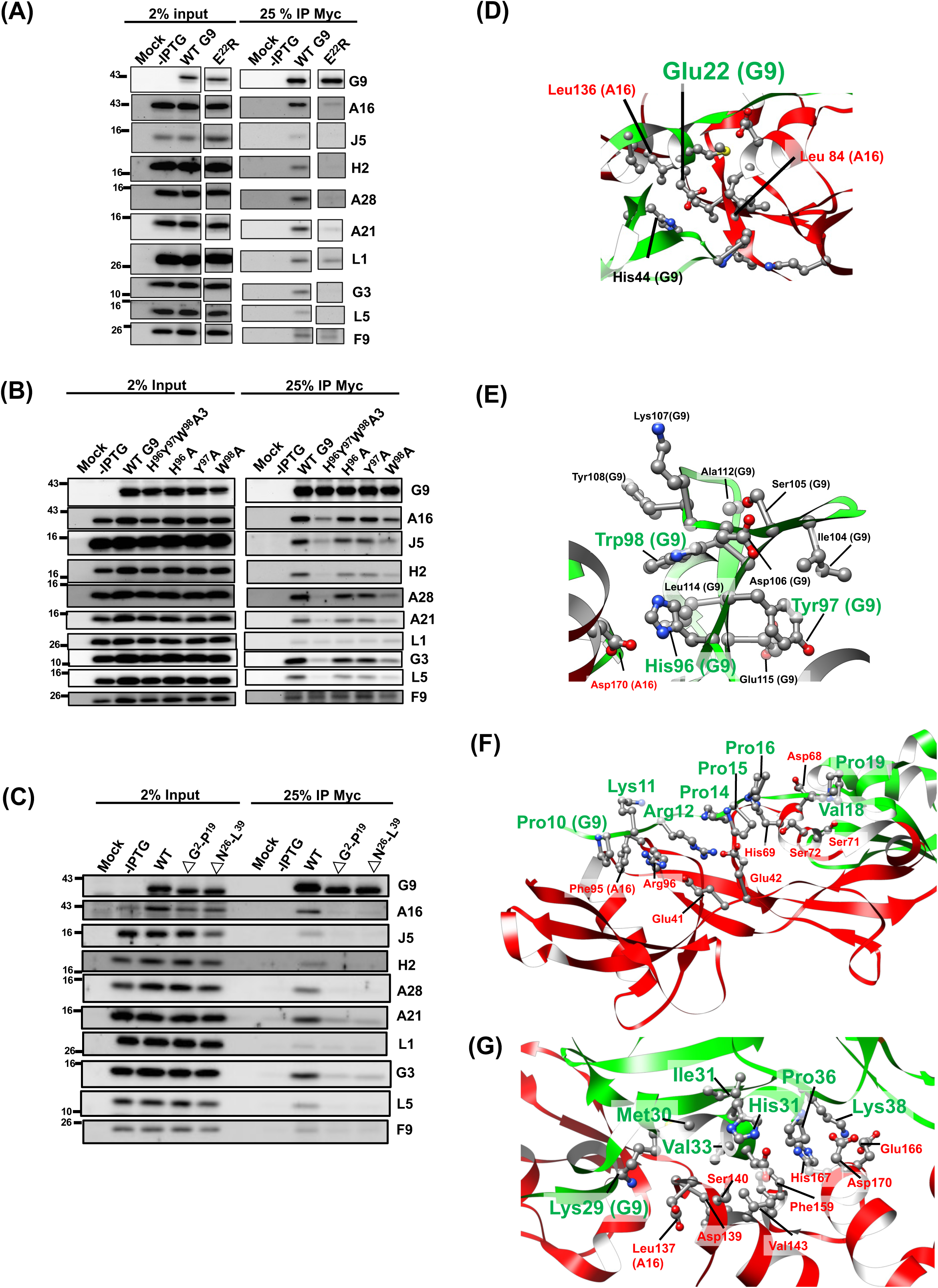
Group 1: G9 Mutants Unable to Bind A16 or Other EFC Components. **(A–C)** Co-immunoprecipitation of lysates from infected/transfected cells expressing G9 mutants. Four mutants (E^22^R (in A) H^96^Y^97^W^98^A3 (in B), ΔG^2^–P^19^ and ΔN^26^–L^39^ (in C)) failed to co-precipitate A16 or other EFC components. (D–G) Structural mapping of key residues based on G9-A16 co-crystal structure (38). Critical residues on G9 (green) and their interacting partners on A16 (red) are highlighted. Cartoon and ball-stick formats depict the spatial relationship of these residues.

### Defective Group 1 G9 mutants disrupted G9-A16 subcomplex and EFC assembly

To investigate the molecular basis of fusion and infectivity defects, we performed co-immunoprecipitation (co-IP) experiments to assess whether G9 mutants interact with A16 protein and other EFC components. The results in Figure 3 showed that the first four G9 mutants (named as the Group 1 mutants), G9^E22R^ (Figure 3A), G9^H96Y97W98A3^ (Figure 3B), G9^ΔG2-P19^ and G9^ΔN26-L39^ (Figure 3C) failed to co-precipitate with A16 as well as other EFC components when the WT G9 protein co-precipitated all EFC components well (Figure 3A-C).

We then looked at the locations of these important residues in the crystal structure of G9-A16 subcomplex(38) (Figure 3D-G) and noticed that E22 is highly conserved in all the G9 orthologues in *Poxviridae* (Figure 1A). Based on the Residue Interaction Network Generator (RING) analyses, E22 forms hydrogen and van der Waals (VDW) interactions with L84 and L136 of A16 protein, respectively (Figure 3D). In addition, E22 is proximal (∼ 3.4 Å) to H44 of G9 protein, suggesting a pH-dependent electrostatic interaction that may stabilize G9 conformation at low pH. The G9^E22R^ mutation likely introduces charge repulsion with H44, disrupting the pH-dependent interaction, implying that the E22-H44 interaction at low pH also contributes to G9 protein function.

The conserved G9 residues H96, Y97, and W98 form a β-strand, in which H96 interacts with D170 of the A16 protein through ionic and van der Waals interactions (Figure 3E). Notably, Y97 and W98 engage in intra-molecular interactions with multiple residues of the G9 protein itself (I104, D106, K107, Y108, A112, L114, E115) via hydrogen bonding, van der Waals forces, and hydrophobic interactions (Figure 3E, Figure S2A). These interactions collectively shape the region into a triple β-stranded structure flanked by a loop (Figure S2B). Structurally, this region serves as a linker between the N-terminal helical domain (residues 1-81) and the C-terminal region (residues 117-340) of G9 (Figure S2B). Consequently, the triple mutant G9^H96Y97W98A3^, which lacks the side chains necessary for these stabilizing interactions, likely disrupts the integrity of this linker domain, leading to structural alterations and functional loss of G9 protein.

In Figure 3F, the deletion mutant G9^ΔG2–P19^ removed an N-terminal loop on G9 which is essential for contacting with the N-terminal region of A16 protein (E41, E42, D68, H69, S71, S72, F95, and R96). Consistent with this structure feature, the G9^ΔG2–P19^ mutant failed to interact with A16 and other EFC components (Figure 3C). Finally, the N26-L39 region of G9 protein was previously shown to be important for superinfection interference by A56 and K2 proteins (41). Here, the co-IP results and the structure interpretation of the G9^ΔN26–L39^ mutant (Figure 3C) showed that this region is important for interactions with multiple residues (L137, D139, S140, V143, F159, E166, H167 and D170) of A16 protein (Figure 3G). Taken together, all the residues defined in the above-mentioned Group 1 mutants are important for G9 and A16 subcomplex interaction as well as the EFC formation.

### Defective Group 2 G9 mutants bound to A16 but failed to form EFC

The Group 2 mutants, G9^G46A^, G9^R78A^, and G9^D120H122D123K126D128A5^ retained A16 binding but failed to co-immunoprecipitate other EFC components (Figure 4A&B). Structurally the mutated residues in this group are located within internal regions of the WT G9 protein and appear to contribute to intra-molecular stabilization. For instance, G46 forms a hydrogen bond with S74, and R78 engages in van der Waals interactions with S116 and Y81 (Figure 4C). Likewise, the clustered mutations in G9^D120H122D123K126D128A5^ disrupt multiple internal contacts, including hydrogen bonding (involving D120, H122, D123, G125, D128, F132), van der Waals interaction (V121 and C127), and ionic interaction (D128 with R155) (Figure 4D). Spatially, both G46 and R78 are located near the N-terminal region of G9 and are not in close contact with A16 (Figure S3A), while the charged cluster (D120H122D123K126D128) lies on the protein surface but remains distal to A16 (Figure S3B). These observations suggest that Group 2 mutations may impair EFC assembly by destabilizing G9’s tertiary structure, while preserving the G9-A16 subcomplex.

**Figure 4.**
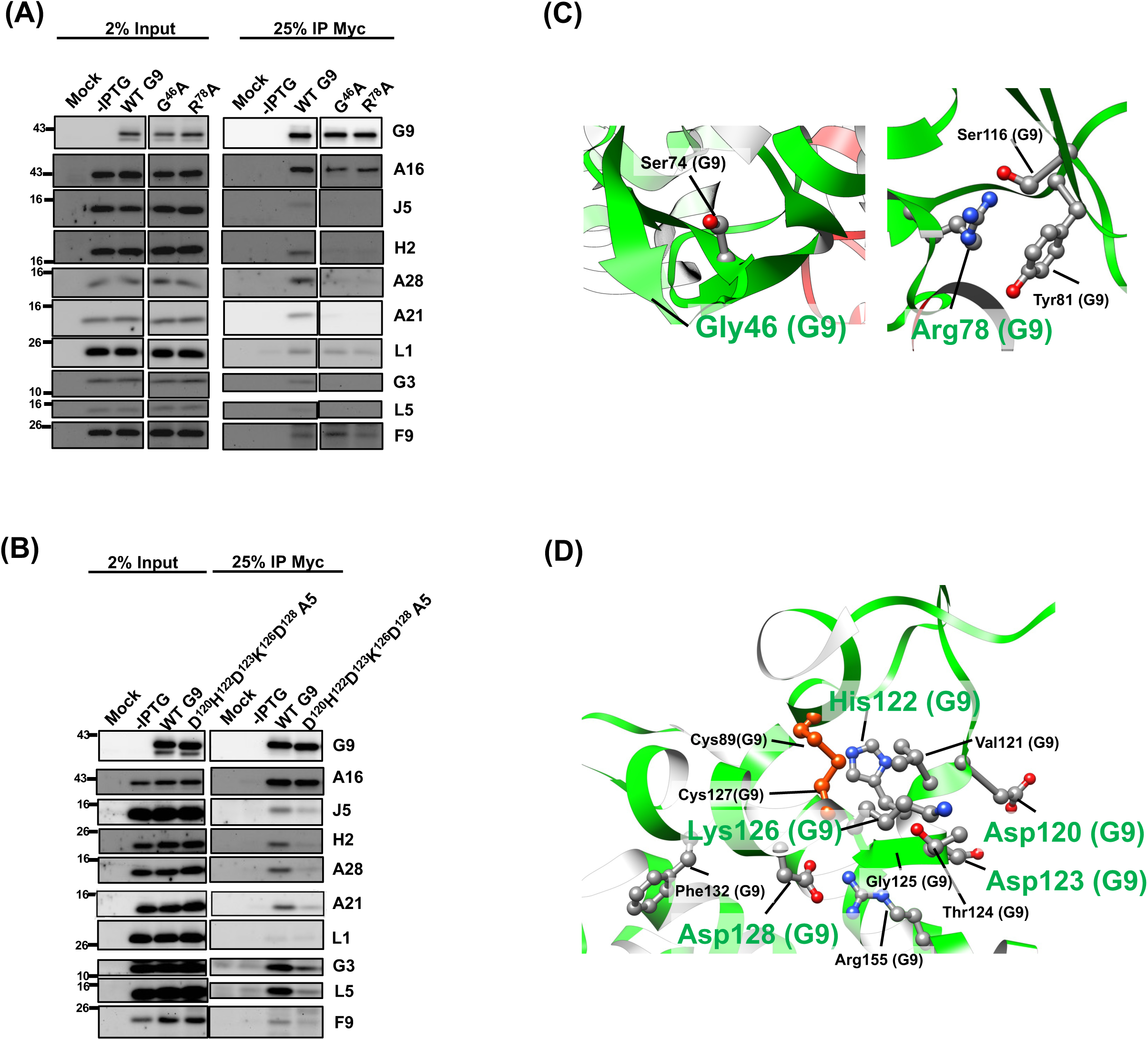
Group 2: G9 Mutants Bound to A16 But Failed to Assemble Full EFC. **(A–B)** Co-IP assays showing G9 mutants (G^46^A, R^78^A (in A) and D^120^H^122^D^123^K^126^D^128^A5 (in B)) retained binding to A16 but failed to associate with other EFC components. **(C–D)** Structural insights into these residues, which are distal to the G9-A16 interface. The critical residues (green) on G9 are involved in intra-G9 interactions important for EFC assembly. The structures were presented in a flat-ribbon cartoon model, and the interaction side-chains were shown in ball and stick format.

### Defective Group 3 G9 mutants, which harbor mutations in the conserved P(R/Y)XCW motif and/or the adjacent loop, retained A16 binding but failed to assemble the EFC

Next, as shown in the Figure 5, the last group of G9 mutants, named as the Group 3 mutants, contained G9^P240R241E242W244A4^ and G9^ΔN273-S290A3^ mutants. Both G9 mutants targeted multiple residues within a region that shared conserved disulfide bonding patterns among vaccinia G9, A16 and J5 proteins (red lines in Figure 5A). Two interesting motifs, the P(R/Y)XCW that surrounded by 4-5 conserved α-helical arrangements (Figure 5B) and three homologous loops linked by disulfide bonds, are in this region (area shaded in gray in Figure 5A). The latter motif was not in the G9-A16 complex crystal structure (38) and the comparable region in A16, G9 and J5 were predicted with Alphafold2 as a closed loop via disulfide bonding (Figure 5C).

**Figure 5.**
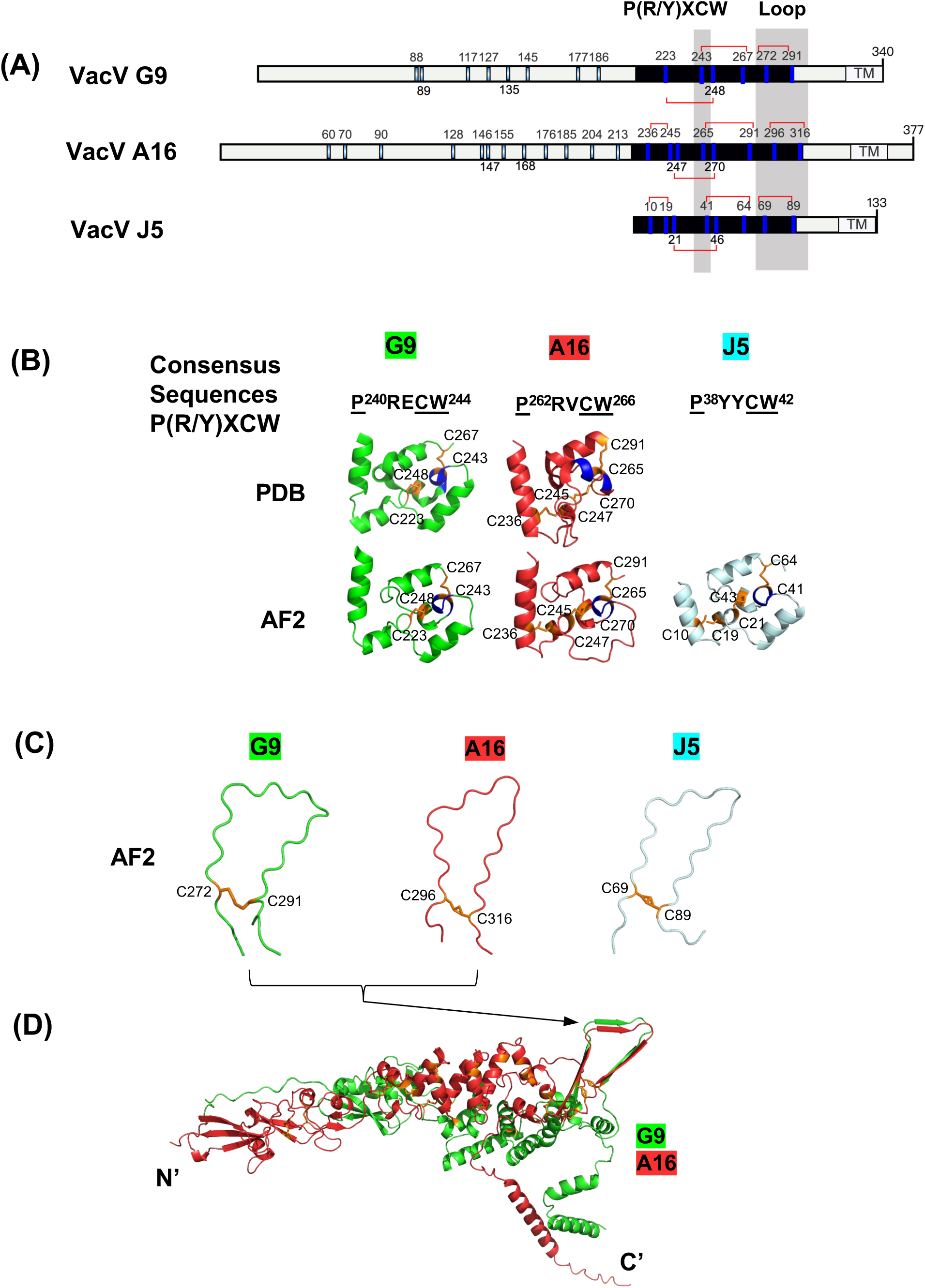
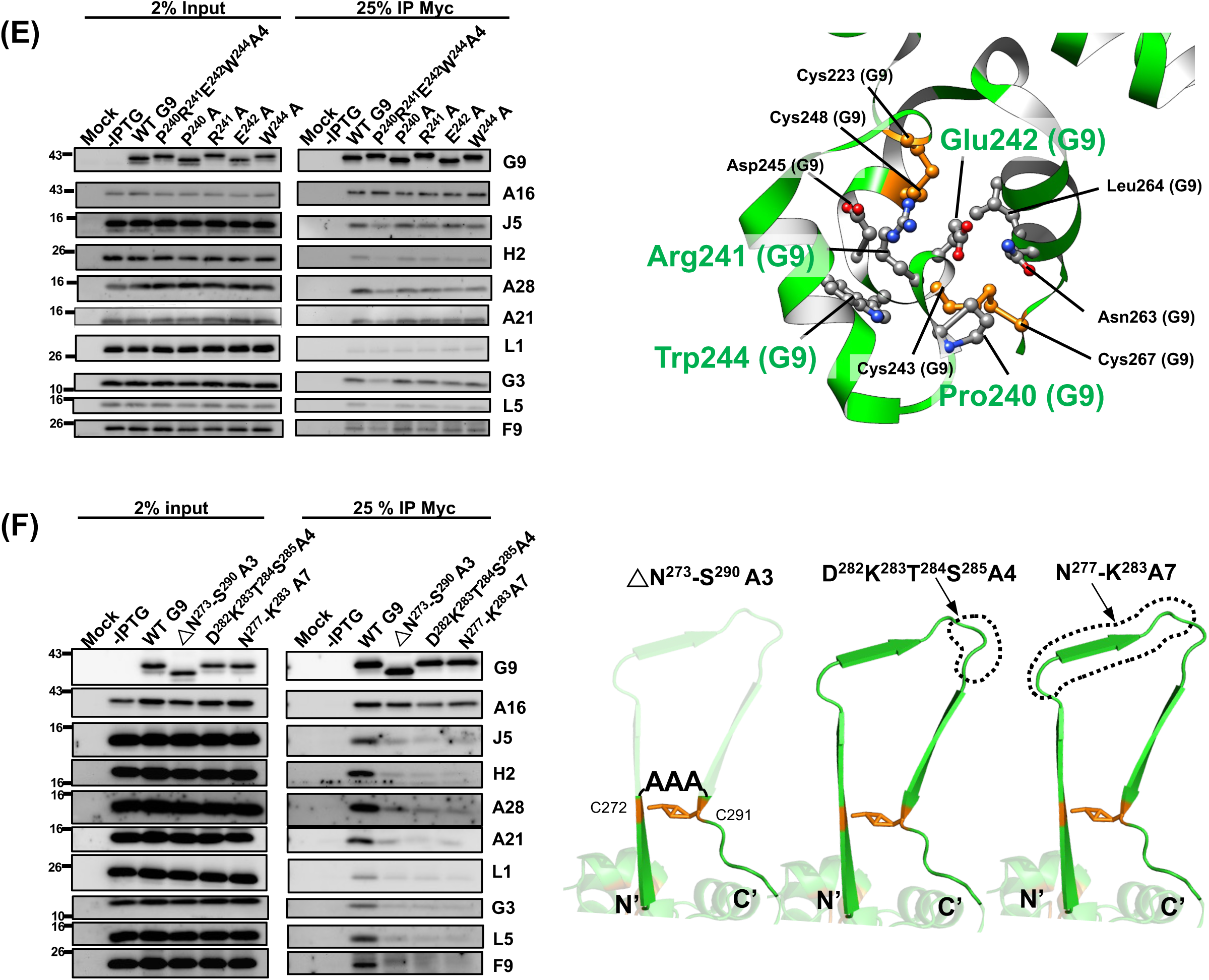
Group 3: Conserved C-terminal G9 Mutants Affected EFC Assembly. **(A)** Schematic representation of vaccinia G9, A16 and J5 protein. Cysteine residues are marked and numbered. The disulfide bonds (red lines) in the conserved P(R/Y)XCW and loop regions (shaded in gray) are shown. The scheme was generated with illustrator of biological sequences (IBS) (42). **(B)** The conserved P(R/Y)XCW motif (the helix marked in blue) in the G9 and A16 crystal structures (PDB) or in the J5 predicted structure from Alphafold2 modeling (AF2) are shown. The conserved disulfide bonds are shown in orange. **(C)** The loop regions formed by disulfide bonds (in orange) in G9, A16, and J5 proteins have not been crystallized so they were individually modeled by AF2, as illustrated. **(D)** The above loops in G9 and A16 appeared as three b-strands when the full length G9-A16 subcomplex was modeled by AF2. **(E)** Co-IP analyses of the tetra-mutant G9 ^P240R241E242W244A4^ and individual mutants G9^P240A^, G9^R241A^, G9^E242A^, G9^W244A^ (the left panel). Structural insights of these target residues (in green) in G9 protein (the right panel). **(F)** Co-IP analyses of three loop mutants, G9_ΔN273–S290A3_, G9_D282K283T284S285A4_ and G9_N277-K283A7_ (the left panel). The mutation designs in the G9 loop region are illustrated (the right panel).

Furthermore, the G9-A16 complex structure covers only truncated forms of the proteins (38). To better understand the interactions within the full-length G9-A16 subcomplex, we employed AlphaFold2 to predict the complete structures in subcomplex form, which predicted that the loops form parallel β-strands at the G9-A16 interface (Figure 5D). We thus constructed two G9 mutants, including a tetra-mutant G9^P240R241E242W244A4^ that eliminated the P(R/Y)XCW and the G9^ΔN273-S290A3^ mutant that deleted the loop.

The co-IP results of the tetra mutant G9^P240R241E242W244A4^ retained the A16 binding but failed to co-IP other EFC components (left panel of Figure 5E). Individual point mutations in this motif (G9^P240A^, G9^R241A^, G9^E242A^, G9^W244A^) showed milder effects, suggesting that multiple side chains cooperate structurally. (left panel of Figure 5E).

Based on G9-A16 complex structure, P240, R241, E242, W244 are not located at the interface of G9-A16 subcomplex and, instead, they formed a compact helix via multiple self-interactions such as H-bonding between R241-W244 and P240- C243; VDW interactions between E242, N263 and L264. The PRECW-containing helix is also surrounded by adjacent three α-helices via interactions with N263 and L264, in addition to the described-above self-interacting residues. Together these intra-molecular interactions facilitate a stable PRECW motif in G9 protein. (right panel of Figure 5E).

Similarly, the G9^ΔN273–S290A3^ mutant, in which the loop spanning residues N273 to S290 was deleted and replaced with three alanine, retained A16 binding but lost interaction with other EFC components (Figure 5F, left panel). To determine which portion of this loop is functionally important, we generated two substitution mutants, G9 ^D282K283T284S285A4^ and G9 ^N277-K283A7^ (Figure 5F). Both mutants exhibited the same phenotype as the G9^ΔN273–S290A3^ mutant, losing virus infectivity but maintaining A16 binding while failing to recruit other EFC proteins. These findings demonstrate that this conserved loop is essential for complete EFC assembly and highlight the critical role of Group 3 residues in stabilizing the fusion-competent architecture of the G9 protein.

## Discussion

In this study, we performed a functional analysis of the vaccinia virus G9 protein and identified residues critical for virus infectivity (Figure 1B), MV-mediated membrane fusion (Figure 2), and Entry Fusion Complex (EFC) formation (Figures 3– 5). We summarized our results in Figure 6A, where residues essential for G9 function are color-coded by severity (red, orange, and yellow). Based on co-immunoprecipitation data, we classified these residues into three functional groups (Figure 6B), which reflect their contributions to G9-A16 subcomplex formation and broader EFC assembly.

**Figure 6.**
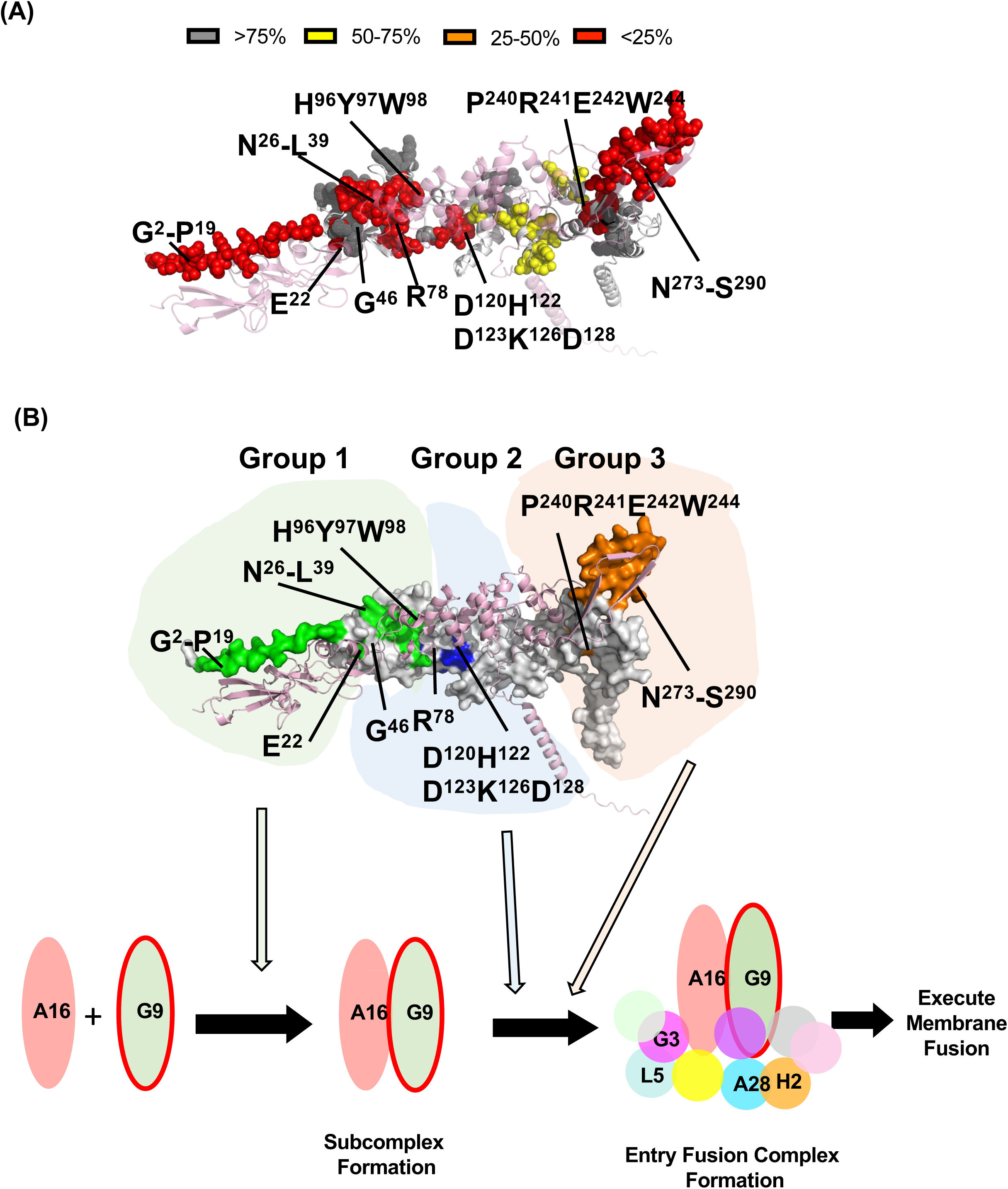
Summary of Functionally Important Regions in vaccinia G9 protein. (**A**). The full-length G9-A16 AF2-predicted model colored with G9 residues that, when mutated, exhibited a significant loss of MV infectivity: mutant titer <25% of WT (in red), ∼25–50% of WT (in orange), ∼50–75% of WT (in yellow) and >75% of WT (in gray). VacV A16 protein is shown in pink. (**B**). The same G9-A16 AF2-predicted model, as shown in A, colored with G9 residues important in Co-IP experiments (green in Group 1, blue in Group2 and orange in Group 3). The overall working model illustrating G9’s Group 1 residues involve in G9-A16 subcomplex formation while Groups 2 and 3 residues facilitates the subsequent EFC formation.

We examined 47 mutations covering approximately 34% of G9 protein sequences. The 9 important G9 mutants are classified into 3 groups. The Group 1 mutants, G9^E22R^, G9^H96Y97W98A3^, G9^ΔG2–P19^, G9^ΔN26–L39^ directly interrupted G9 and A16 protein-protein interaction and as well as EFC formation (green area, Figure 6B). The Group 2 mutant G9 proteins, including G9^G46A^, G9^R78A^, and the G9^D120H122D123K126D128A5^, bound to A16 protein but not to other EFC components (blue area, Figure 6B). Finally, the Group 3 G9^P240R241E242W244A4^ and G9 ^△N273-S290A3^ mutants are focused on two motifs that are conserved among G9, A16, and J5. Group 3 mutations, similar to Group 2, bound to A16 protein but not to other EFC components (orange, Figure 6B). Both Group 2 and 3 mutations are far from the G9 and A16 interaction interphase, explaining why these mutations did not affect G9/A16 subcomplex formation. However, it remains unclear how these Group 2 and 3 mutations interrupted the EFC assembly and it will require future investigation. At this point, our data implied that G9 and A16 subcomplex formation may be pre-request for the viral EFC assembly (bottom panel of Figure 6B).

Interestingly, in Group 3, we identified a functionally important motif, P(R/Y)XCW, within a conserved disulfide bonding region among vaccinia G9, A16, and J5 proteins (Figure 5A&B). Sequence alignment of orthologues of G9, A16, and J5 in 28 *Poxviridae* members (e.g., VACV, CMLV, CPXV, ECTV, MPXV, etc.) together revealed a highly conserved consensus sequence in this region (Fig. S4A). We previously showed that vaccinia G9 homologues are present in members of *Nucleocytoviricota* (40). We expanded this analysis and found that these G9 homologs in giant viruses also share this motif (Figure S4B). These findings suggest that this conserved domain is a fundamental element of viral membrane fusion machinery and may have been evolutionarily retained among large DNA viruses.

The identification of critical residues in vaccinia G9 protein delineates distinct structural and functional roles within the EFC. Group 1 residues are indispensable for forming the G9-A16 subcomplex and serve as scaffolding elements for EFC nucleation. Group 2 and 3 residues, although not required for A16 interaction, are crucial for downstream complex assembly. Collectively, our results reveal that G9 contains multiple discrete domains that mediate sequential steps of EFC formation and membrane fusion. By identifying conserved and structurally vital residues, this study deepens our understanding of poxvirus EFC and identifies potential targets for antiviral strategies.

## Acknowledgments

We thank Bernard Moss for providing the IPTG-inducible virus vG9Li-HA. We are also grateful to Sue-Ping Lee, Yae-Huei Liou, and Wen-Li Pon at the IMB Imaging Core, Academia Sinica, for their support with imaging data analysis, and to Hsin-Nan Lin at the IMB Bioinformatics Core, Academia Sinica, for assistance with bioinformatic analyses.

## Supplemental figure legends

**Fig. S1. Normal MV morphogenesis of G9 mutant viruses.**>

BSC40 cells were infected with vG9Li-HA (5 PFU/cell, 1 h at 37 °C), followed by transfection with either wild-type or mutant G9 plasmids. Infected/transfected cells were incubated in complete medium without IPTG for 24 h and processed for transmission electron microscopy as described in *Materials and Methods*. Scale bar, 1 µm.

**Fig.S2. A conserved H96Y97W98 segment forms a linker-like structure in G9.**

(A) Conserved residues H96Y97W98 (green) interact with nearby residues (I104, D106, K107, Y108, A112, L114; orange) via hydrogen bonds, van der Waals, and hydrophobic contacts.

(B) The triple β-strand motif likely acts as a linker domain bridging the N-terminal helical domain (aa 1–81) and the C-terminal region (aa 117–340) of G9.

**Fig. S3. Group 2 G9 mutations map to regions distant from A16.**

(A) G46 and R78 (green) form internal contacts with S74, Y81, and S116 (orange) within G9 (gray cartoon), but do not directly contact A16 (pink cartoon) in the G9-A16 complex.

(B) The surface cluster D120, H122, D123, K126, and D128 (green) interact with adjacent residues V121, G125, C127, F132, and R155 (orange) but remain spatially distant from A16.

**Fig. S4. The P(R/Y)XCW motif is conserved across Poxviridae and Nucleocytoviricota.**

(A) Sequence logo generated from 28 Poxviridae members members (described in (35)) shows high conservation of the P(R/Y)XCW motif. (B) Alignment of 11 G9 homologous genes from five Nucleocytoviricota members (Klosneuvirus, Frog virus 3, Tupanvirus deep ocean, Medusavirus, Pacmanvirus A23) also reveals a conserved motif. Sequences were aligned using MacVector (v16.0.10), and consensus logos were generated via WebLogo3 (http://weblogo.threeplusone.com/).

## Reference

1. Moss B. 1996. Poxviridae: the viruses and thier replication. Fields virology:2637–2671.

2. Goebel SJ, Johnson GP, Perkus ME, Davis SW, Winslow JP, Paoletti E. 1990. The complete DNA sequence of vaccinia virus. Virology 179:247–266.

3. Maruri-Avidal L, Weisberg AS, Moss B. 2013. Direct formation of vaccinia virus membranes from the endoplasmic reticulum in the absence of the newly characterized L2-interacting protein A30.5. J Virol 87:12313–26.

4. Chlanda P, Carbajal MA, Cyrklaff M, Griffiths G, Krijnse-Locker J. 2009. Membrane rupture generates single open membrane sheets during vaccinia virus assembly. Cell host & microbe 6:81–90.

5. Schmelz M, Sodeik B, Ericsson M, Wolffe EJ, Shida H, Hiller G, Griffiths G. 1994. Assembly of vaccinia virus: the second wrapping cisterna is derived from the trans Golgi network. J Virol 68:130–47.

6. Smith GL, Vanderplasschen A. 1998. Extracellular enveloped vaccinia virus: entry, egress, and evasion. Coronaviruses and Arteriviruses:395–414.

7. Ward BM, Moss B. 2001. Vaccinia virus intracellular movement is associated with microtubules and independent of actin tails. J Virol 75:11651–63.

8. Lynn H, Howell LM, Diefenbach RJ, Newsome TP. 2018. Phototracking Vaccinia Virus Transport Reveals Dynamics of Cytoplasmic Dispersal and a Requirement for A36R and F12L for Exit from the Site of Wrapping. Viruses 10.

9. Hollinshead M, Rodger G, Van Eijl H, Law M, Hollinshead R, Vaux DJ, Smith GL. 2001. Vaccinia virus utilizes microtubules for movement to the cell surface. J Cell Biol 154:389–402.

10. van Eijl H, Hollinshead M, Rodger G, Zhang W-H, Smith GL. 2002. The vaccinia virus F12L protein is associated with intracellular enveloped virus particles and is required for their egress to the cell surface. Journal of General Virology 83:195–207.

11. Lin CL, Chung CS, Heine HG, Chang W. 2000. Vaccinia virus envelope H3L protein binds to cell surface heparan sulfate and is important for intracellular mature virion morphogenesis and virus infection in vitro and in vivo. J Virol 74:3353–65.

12. Hsiao JC, Chung CS, Chang W. 1999. Vaccinia virus envelope D8L protein binds to cell surface chondroitin sulfate and mediates the adsorption of intracellular mature virions to cells. J Virol 73:8750–61.

13. Chung CS, Hsiao JC, Chang YS, Chang W. 1998. A27L protein mediates vaccinia virus interaction with cell surface heparan sulfate. J Virol 72:1577–85.

14. Chiu WL, Lin CL, Yang MH, Tzou DL, Chang W. 2007. Vaccinia virus 4c (A26L) protein on intracellular mature virus binds to the extracellular cellular matrix laminin. J Virol 81:2149–57.

15. Ojeda S, Domi A, Moss B. 2006. Vaccinia virus G9 protein is an essential component of the poxvirus entry-fusion complex. J Virol 80:9822–30.

16. Ojeda S, Senkevich TG, Moss B. 2006. Entry of vaccinia virus and cell-cell fusion require a highly conserved cysteine-rich membrane protein encoded by the A16L gene. J Virol 80:51–61.

17. Senkevich TG, Ojeda S, Townsley A, Nelson GE, Moss B. 2005. Poxvirus multiprotein entry–fusion complex. Proceedings of the National Academy of Sciences 102:18572–18577.

18. Townsley AC, Senkevich TG, Moss B. 2005. Vaccinia virus A21 virion membrane protein is required for cell entry and fusion. J Virol 79:9458–69.

19. Senkevich TG, Ward BM, Moss B. 2004. Vaccinia virus A28L gene encodes an essential protein component of the virion membrane with intramolecular disulfide bonds formed by the viral cytoplasmic redox pathway. J Virol 78:2348–56.

20. Senkevich TG, Moss B. 2005. Vaccinia virus H2 protein is an essential component of a complex involved in virus entry and cell-cell fusion. J Virol 79:4744–54.

21. Izmailyan RA, Huang CY, Mohammad S, Isaacs SN, Chang W. 2006. The envelope G3L protein is essential for entry of vaccinia virus into host cells. J Virol 80:8402–10.

22. Townsley AC, Senkevich TG, Moss B. 2005. The product of the vaccinia virus L5R gene is a fourth membrane protein encoded by all poxviruses that is required for cell entry and cell-cell fusion. J Virol 79:10988–98.

23. Brown E, Senkevich TG, Moss B. 2006. Vaccinia virus F9 virion membrane protein is required for entry but not virus assembly, in contrast to the related L1 protein. J Virol 80:9455–64.

24. Bisht H, Weisberg AS, Moss B. 2008. Vaccinia virus l1 protein is required for cell entry and membrane fusion. J Virol 82:8687–94.

25. Satheshkumar PS, Moss B. 2009. Characterization of a newly identified 35-amino-acid component of the vaccinia virus entry/fusion complex conserved in all chordopoxviruses. J Virol 83:12822–32.

26. Moss B. 2012. Poxvirus Cell Entry: How Many Proteins Does it Take? Viruses 4:688–707.

27. Wagenaar TR, Ojeda S, Moss B. 2008. Vaccinia virus A56/K2 fusion regulatory protein interacts with the A16 and G9 subunits of the entry fusion complex. J Virol 82:5153–60.

28. Nelson GE, Wagenaar TR, Moss B. 2008. A conserved sequence within the H2 subunit of the vaccinia virus entry/fusion complex is important for interaction with the A28 subunit and infectivity. Journal of virology 82:6244–6250.

29. Wolfe CL, Moss B. 2011. Interaction between the G3 and L5 proteins of the vaccinia virus entry–fusion complex. Virology 412:278–283.

30. Hong G-C, Tsai C-H, Chang W, Shisler JL. 2020. Experimental Evolution To Isolate Vaccinia Virus Adaptive G9 Mutants That Overcome Membrane Fusion Inhibition via the Vaccinia Virus A56/K2 Protein Complex. Journal of Virology 94:e00093–20.

31. Laliberte JP, Weisberg AS, Moss B. 2011. The membrane fusion step of vaccinia virus entry is cooperatively mediated by multiple viral proteins and host cell components. PLoS pathogens 7:e1002446.

32. Lin S, Yue D, Yang F, Chen Z, He B, Cao Y, Dong H, Li J, Zhao Q, Lu G. 2023. Crystal structure of vaccinia virus G3/L5 sub-complex reveals a novel fold with extended inter-molecule interactions conserved among orthopoxviruses. Emerging Microbes & Infections 12:e2160661.

33. Su H-P, Garman SC, Allison TJ, Fogg C, Moss B, Garboczi DN. 2005. The 1.51-Å structure of the poxvirus L1 protein, a target of potent neutralizing antibodies. Proceedings of the National Academy of Sciences 102:4240–4245.

34. Diesterbeck US, Gittis AG, Garboczi DN, Moss B. 2018. The 2.1 Å structure of protein F9 and its comparison to L1, two components of the conserved poxvirus entry-fusion complex. Scientific Reports 8:16807.

35. Kao C-F, Liu C-Y, Hsieh C-L, Carillo KJD, Tzou D-LM, Wang H-C, Chang W. 2023. Structural and functional analyses of viral H2 protein of the vaccinia virus entry fusion complex. Journal of Virology 0:e01343–23.

36. Kao C-F, Tsai M-H, Carillo KJ, Tzou D-L, Chang W. 2023. Structural and functional analysis of vaccinia viral fusion complex component protein A28 through NMR and molecular dynamic simulations. PLOS Pathogens 19:e1011500.

37. Diesterbeck U, Tak, A., Gittis, A.G., Garboczi, D.N., Moss, B. 2024. The crystal structure of protein A21, a component of the conserved poxvirus entry-fusion complex.

38. Yang F, Lin S, Chen Z, Yue D, Yang M, He B, Cao Y, Dong H, Li J, Zhao Q. 2023. Structural basis of poxvirus A16/G9 binding for sub-complex formation. Emerging Microbes & Infections 12:2179351.

39. Chang HW, Yang CH, Luo YC, Su BG, Cheng HY, Tung SY, Carillo KJD, Liao YT, Tzou DM, Wang HC, Chang W. 2019. Vaccinia viral A26 protein is a fusion suppressor of mature virus and triggers membrane fusion through conformational change at low pH. PLoS Pathog 15:e1007826.

40. Kao S, Kao C-F, Chang W, Ku C. 2023. Widespread distribution and evolution of poxviral entry-fusion complex proteins in giant viruses. Microbiology Spectrum 11:e04944–22.

41. Cotter CA, Moss B. 2020. Mutations near the N terminus of vaccinia virus G9 protein overcome restrictions on cell entry and syncytium formation imposed by the A56/K2 fusion regulatory complex. Journal of Virology 94:10.1128/jvi. 00077-20.

42. Xie Y, Li H, Luo X, Li H, Gao Q, Zhang L, Teng Y, Zhao Q, Zuo Z, Ren J. 2022. IBS 2.0: an upgraded illustrator for the visualization of biological sequences. Nucleic Acids Research 50:W420–W426.

